# Towards Longitudinal Characterization of Multiple Sclerosis Atrophy Employing SynthSeg Framework and Normative Modeling

**DOI:** 10.1101/2024.09.17.613272

**Authors:** Pedro M. Gordaliza, Nataliia Molchanova, Maxence Wynen, Pietro Maggi, Joost Janssen, Jaume Banus, Alessandro Cagol, Cristina Graziera, Meritxell Bach Cuadra

**Affiliations:** CIBM Center for Biomedical Imaging, Switzerland; Department of Radiology, Lausanne University Hospital (CHUV) and University of Lausanne (UNIL), Lausanne, Switzerland; University of Applied Sciences Western Switzerland (HES-SO), Switzerland; ICTEAM, Universite catholique de Louvain, Louvain-la-Neuve, Belgium; Neuroinflammation Imaging Lab (NIL), Universite catholique de Louvain, Belgium; Cliniques universitaires Saint-Luc, Universite catholique de Louvain, Belgium; Department of Child and Adolescent Psychiatry, Institute of Psychiatry (HGUGM-IiSGM) and CIBERSAM, Instituto de Salud Carlos III, Madrid, Spain; Translational Imaging in Neurology (ThINk), Department of Medicine and Biomedical Engineering, Basel Hospital and University of Basel (USB), Switzerland

## Abstract

Multiple Sclerosis (MS) is a complex neurodegenerative disease characterized by heterogeneous progression patterns. Traditional clinical measures like the Expanded Disability Status Scale (EDSS) inadequately capture the full spectrum of disease progression, highlighting the need for advanced Disease Progression Modeling (DPM) approaches. This study harnesses cutting-edge neuroimaging and deep learning techniques to investigate deviations in subcortical volumes in MS patients. We analyze T1-weighted and Fluid-attenuated inversion recovery (FLAIR) Magnetic Resonance Imaging (MRI) data using advanced DL segmentation models, *SynthSeg*^+^ and *SynthSeg-WMH*, which address the challenges of conventional methods in the presence of white matter lesions. By comparing subcortical volumes of 326 MS patients to a normative model from 37,407 healthy individuals, we identify significant deviations that enhance our understanding of MS progression. This study highlights the potential of integrating DL with normative modeling to refine MS progression characterization, automate informative MRI contrasts, and contribute to data-driven DPM in neurodegenerative diseases.

## 1 Introduction

Multiple Sclerosis (MS) is a chronic autoimmune neurodegenerative disease affecting approximately 2.3 million people globally [14]. MS presents significant challenges due to its heterogeneous manifestations and unclear etiology [21]. Traditionally, MS is clinically characterized by the pseudo-quantitative Expanded Disability Status Scale (EDSS), which pseudo-quantitatively measures the level of disability [42] and, to a limited extent, tracks cognitive impairment over time [4, 41]. However, this approach represents an insufficient proxy for essential Disease Progression Modeling (DPM), unable to characterize the set of underlying mechanisms such as neuroinflammation and degenerative processes recently known as Smouldering MS [19, 34].

Magnetic Resonance Imaging (MRI) plays a crucial role in the clinical assessment of MS patients, for diagnosis, prognosis, and Disease Modifying Therapies (DMT) assessment [42]. Structural MRI provides information on focal lesion dissemination in the brain and spinal cord, as well as brain structure characterization [43]. While cross-sectional analysis of Fluid-Attenuated Inversion Recovery (FLAIR) and T1-weighted (T1w) contrasts are routinely acquired in clinical settings for lesion identification [28], lesion-derived biomarkers have shown controversial or limited relationships with disease phenotypes and progression. This highlights the need for more robust approaches to capture disease progression over time, particularly for DPM.

In contrast, brain structure evolution assessment provides atrophy-related imaging biomarkers of well-proven neurodegeneration, which is more acute in MS patients [17, 22]. Studies employing global and regional atrophy imaging biomarkers have associated deviations from control measures for MS patients with cognitive impairment [5, 13], depression [3], and physical disability [23, 32].

The literature shows how subcortical structures are implicated in the pathophysiology of MS due to their involvement in key neurological functions often compromised in the disease, compared with cortical regions, which are more difficult to analyze in conventional MRI sequences [33]. Findings across several studies have revealed that gray matter atrophy in MS is more pronounced, particularly in subcortical regions such as the thalamus [2] and the putamen [27], compared to the rates of atrophy in healthy controls [43]. This regional Grey Matter (GM) analysis enables deeper studies about MS mechanisms and allows further disease progression modeling [5]. Recent models to characterize the longitudinal trajectory of brain region volumes for MS patients have been published [10, 26], emphasizing the importance of longitudinal data projection in understanding disease progression.

However, these approaches face several challenges: they are difficult to implement in clinical practice, may be limited by small sample sizes, and struggle to account for the heterogeneity of MS evolution. Furthermore, they do not provide a direct comparison to healthy population norms. To address these limitations, we propose to measure the deviance of each patient at each time-point concerning a longitudinal trajectory model based on thousands of healthy subjects. This approach has been successfully applied to other neurological diseases through the normative modeling framework [25, 39], allowing for a more robust and clinically applicable method of characterizing individual disease trajectories.

Practically, manual segmentation of main brain structures for thousands of subjects is unfeasible. Models often rely on automatic segmentation provided by tools such as FreeSurfer [15] or more recently DL approaches [9], which provide acceptable segmentations for quality T1w images of healthy subjects. However, these tools tend to be more uncertain on volume estimations for MS patients who present white matter lesions (WML), where more reliable information is usually contained within the FLAIR contrast (see Fig.1) [35]. Recent advancements in DL-based segmentation algorithms have improved on this issue. SynthSeg [7] allows for obtaining reliable volumetric measures when employing FLAIR [36], and its new version, *SynthSeg-WMH* [29], can handle the presence of WML. These developments could accelerate the use of regional atrophy-related imaging biomarkers leveraging normative modeling fed by regions segmented through DL models, empowering speed and reliability.

To explore this potential and address the limitations of previous approaches, we present a pilot study investigating deviations in brain morphometry with three key innovations: 1) utilizing a normative *healthy* brain model rather than MS-specific models [26], 2) comparing segmentations based on both T1w and FLAIR images, which is rarely done in MS studies [36], and 3) employing a tissue segmentation algorithm that jointly segment subcortical structures and white matter lesions (WML) rather than first finding lesions and lesion-filling the input images. Our approach leverages the most advanced deep learning (DL) tools for domain agnostic segmentation of subcortical volumes from MR images (FLAIR and T1w) and utilizes the CentileBrain normative model, based on 37,407 healthy individuals [18]. This methodological combination allows for a more comprehensive and potentially more accurate assessment of brain morphometry in MS patients compared to traditional T1w-only approaches that do not account for WML or utilize normative modeling.

## 2 Materials and Methods

### 2.1 Materials

We analyzed a heterogeneous dataset comprising 326 MS patients from five sources: three in-house datasets (Lausanne University Hospital (CHUV), Louvain Neuroinflammation Imaging Lab (NIL), and Imaging Axonal Damage & Repair in Multiple Sclerosis (INsIDER) [20]) and two public datasets (MICCAI-MSSEG2016 challenge [8] and OpenMSLong [30]). All patients underwent Fluid Attenuated Inversion Recovery (FLAIR) MRI scanning and T1w contrasts (i.e., Magnetization Prepared RApid Gradient Echo -MPRAGE-or MP2RAGE). Imaging protocols varied across datasets, with magnetic field strengths ranging from 1.5T to 3T. For OpenMSLong, Anonymous Dataset 1, and Anonymous Dataset3, two time points were available for some subjects, resulting in a total of 460 3D-FLAIR and T1w MRI scans (Table 1).

**Table 1.**
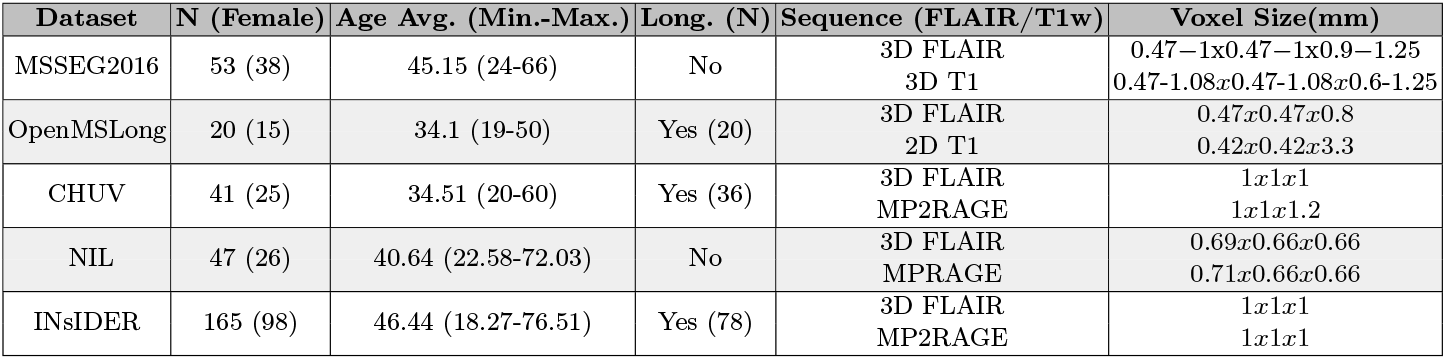
Summary of available datasets.

### 2.2 Methods

In this study, we developed a processing framework for the automatic segmentation of subcortical regions in MS patients using T1w and FLAIR contrasts and the evaluation of deviations from the expected volumes from a large-scale healthy brain model. As depicted in Fig.2, each MRI contrast independently feeds into two different *SynthSeg* models. These models are used to obtain the surrogate ground truth for each patient (*i*), contrast (*c*), subcortical region (*r*), algorithm (*a*), and sex (*s*) denoted as *y*_*icras*_. The *CentileBrain* model [18] employs patient-specific age and sex covariates to estimate the subcortical volume of a healthy subject of such age and sex, represented as *ŷ*_*irs*_. Finally, *y*_*icra*_ and *ŷ*_*irs*_ are used to extract evaluation metrics to assess the deviation from normative values.

#### Subcortical Volumes Segmentation

Automatic segmentation of subcortical structures is particularly challenging in MS patients due to the presence of focal and sparse MS lesions. Tools such as the FreeSurfer suite [15] tend to be less reliable as they depend more on T1w images, which are less informative than FLAIR sequences for depicting MS lesions [35, 36]. In this work, we employ segmentation methods designed with a resolution and contrast-agnostic approach [7]. We perform the segmentation of all subcortical regions using both T1w images and FLAIR sequences, as well as the DL algorithms *SynthSeg*^+^ [7] and *SynthSeg-WMH* [29], referred to as *y*_*icra*_. *SynthSeg*^+^ is a more robust version of the previously released *SynthSeg* [6], which accounts for image quality beyond resolution and contrast agnosticism. *SynthSeg-WMH* is a branch of the former, intended for more reliable segmentation in the presence of WML. Note that the most robust version of *SynthSeg*^+^ is optional and was enabled for this study; no extra parameter setting is needed for any of the DL segmentation algorithms.

#### Subcortical Volumes Site Harmonization

Beyond the mentioned confounders, age and sex, multi-site MRI studies face the challenge of variability in imaging protocols, which can introduce site-related biases in the data. To address this, we used ComBat-GAM harmonization [16], an advanced technique that combines the ComBat method with Generalized Additive Models (GAMs). ComBat-GAM effectively harmonizes neuroimaging data by accounting for age and sex, in addition to mitigating site effects, making it suitable for studies involving large, heterogeneous datasets. By applying ComBat-GAM, we ensure that the sub-cortical volume estimates, *y*_*icras*_, derived from our segmentation methods are comparable across different sites, thereby enhancing the reliability of our normative models and subsequent analyses.

#### Normative modeling

A brain normative model is a statistical framework that characterizes the healthy range of brain structure across a large population by leveraging longitudinal data [31, 39], allowing for the disentanglement of normal changes from deviations linked to neurological diseases. The power of a normative model resides in the sample size used for training, which helps to characterize biological variability across the lifespan better, accounting for confounders of brain changes beyond age, such as sex, and robust model selection. Recently, the *CentileBrain* [18] model trained with 37,407 healthy individuals (53.3% females, age range 3-90) from 81 datasets has been released. The *CentileBrain* employs Multivariate Factorial Polynomial Regression (MFPR) [38] algorithm which performs robustly on healthy subjects, generating sex-specific normative models for the volume estimation of subcortical regions in both right and left brain hemispheres: Thalamus, Caudate, Putamen, Pallidum, Hippocampus, Amygdala, and Accumbens. The same regions are given by both DL segmentation models, making *CentileBrain* ideal for our purposes since subcortical degeneration has been found to be more significant in MS patients [5, 43].

#### Evaluation

Our evaluation process consisted of several steps first to assess which surrogate truths *y*_*icras*_ coincide with expected findings as well as those given from the normative deviation, and to quantify the deviation of MS patients from the normative brain volume trajectories. Notably, while *SynthSeg-WMH* is theoretically more suitable for MS lesion-rich environments, its recent development necessitates comparison with its more validated counterpart, *SynthSeg*^+^, to rigorously assess performance and reliability in this specific context.

### 1. Preliminary comparison with literature values

To ensure our volume estimations are within a reasonable range, we conducted a preliminary comparison with values reported in the literature. We compared our average subcortical volume estimations from both T1w and FLAIR-based segmentations (using both SynthSeg models) against values from studies that performed a brief review of automatic segmentation on T1w images from several classic segmentation software tools, as described in [37]. This comparison serves as an initial validation step, acknowledging that exact matches are not expected due to methodological differences, particularly with our novel FLAIR-based approach. Potential biases and implications of this comparison will be addressed in the discussion section. For each subcortical region, hemisphere, contrast, and algorithm, the Kolmogorov-Smirnov (K-S) test was used to compute p-values that quantify the similarity between our segmented volume distributions and the reference values provided in the literature for MS patients. A higher p-value indicates a closer match to the literature-reported mean values. These p-values were then used to identify and highlight the best matching segmentation approaches, ensuring the accuracy and reliability of our methods.

### 2. Subcortical Volume Longitudinal Trajectory Analysis

MFPR was applied to analyze the age-related trajectories of the harmonized subcortical volumes, *y*_*icras*_, stratified by sex and the tissue segmentation model employed. For a more comprehensive understanding of these trajectories, the Root Mean Square Error (RMSE) between the real volumes (*y*_*icras*_) and the estimated healthy vol-umes (*ŷ*_*irs*_) was calculated across all patients for each region: 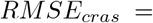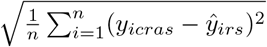 where *n* is the number of scans. This *RMSE*_*cras*_ provides an estimation of the average deviation of the MS cohort from the normative model predictions, stratified by sex, contrast, DL model, and subcortical region.

### 3. Individual Deviation Scores and Longitudinal Deviation

For each patient and each region, we computed a Z-score to quantify the individual deviation from the normative value: 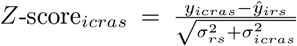 Note that the uncertainty normalization term, 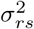, which assumes normality, is approximated using the reported RMSE model (see supplement in [18]) as in [11,12,24]. Finally, from the set of individual scores, we approximated the *longitudinal deviation trajectory*. We plotted the Z-scores against age for each region and hemisphere to visualize the longitudinal trajectories of deviations. Regression lines were fitted to these plots to identify trends in deviations across the lifespan.

## 3 Results

### On the Robustness and Reliability of Subcortical Segmentations

Figure 3 shows the average volume per region and hemisphere for each of the contrast (FLAIR and T1w) and segmentation model (*SynthSeg*^+^ and *SyntheSeg-WMH*) combinations, along with comparisons to reference values for each subcortical volume in the literature [37] (depicted as horizontal lines in Fig.3). Comparing the four combinations, we found that for both hemispheres of the caudate, pallidum, and putamen, the four options for volume extraction did not present significant differences (ANOVA, p > 0.05). In contrast, for the amygdala, accumbens, hippocampus, and thalamus, differences were primarily due to the DL model used, except for the thalamus, where differences were also observed between using FLAIR or T1w (e.g. average left-thalamus volume employing FLAIR and *SynthSeg*^+^ is 7492.2 *±*112.1 vs 6248*±* 101.4 when employing T1w and *SynthSeh-WMH*). Regarding the comparison with reference values extracted from the literature using the K-S test, it was observed that in regions where significant differences existed between the segmentation pipelines, the extraction model was consistent between hemispheres. Segmentations using *SynthSeg-WMH* were closer to the reference values (e.g., 600 *±*100 for left accumbens), with no significant differences between the contrasts used, except for the thalamus volume. The thalamus volume showed greater similarity to the reference value when segmented using the FLAIR sequence and *SynthSeg*^+^ (*p − val* = 0.548).

**Fig 1.**
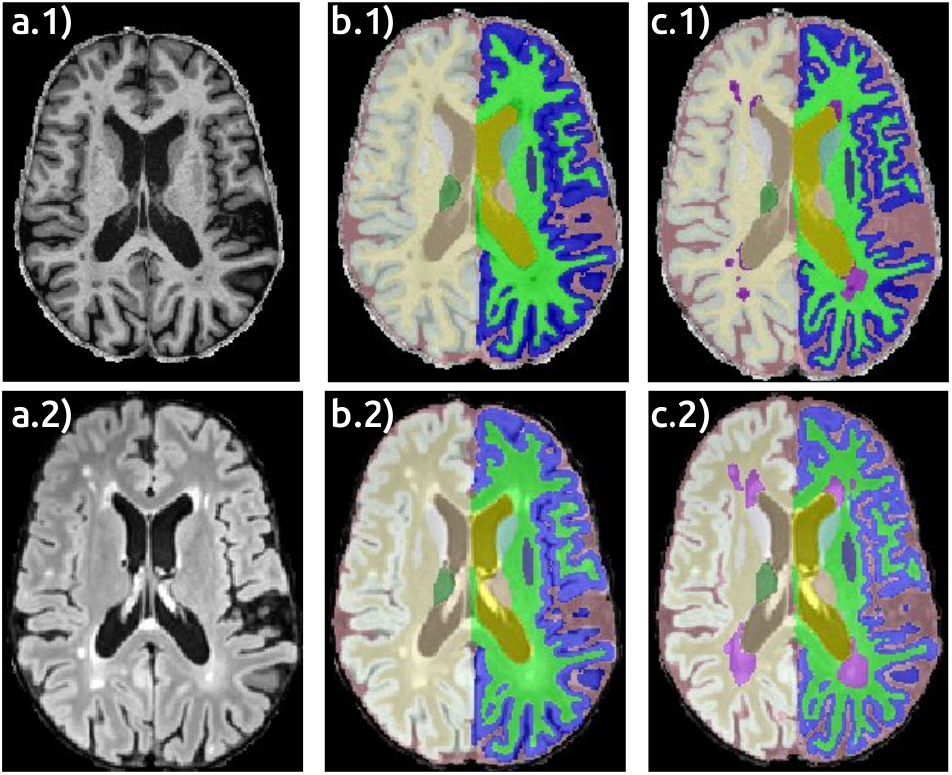
The a) column shows an axial slice of a T1-weighted (T1) image and a Fluid-Attenuated Inversion Recovery (FLAIR) image, respectively. The b) column contains the corresponding *SynthSeg*^+^ [7] segmentation, while the c) column depicts the *SynthSeg-WMH* [29] segmentation. Note the White Matter Lesions (WML) in violet.

**Fig 2.**
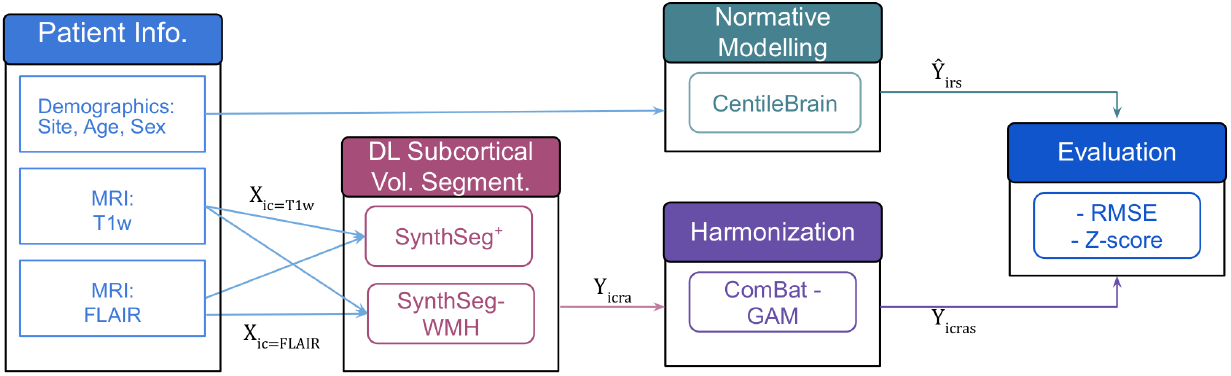
Pipeline: Each available MRI contrast, *Xic* (where *i* represents an MS patient and *c* the contrast, either T1w or FLAIR), independently feeds both *SynthSeg* models (*a*): *SynthSeg*^+^ [7] and *SynthSeg-WMH* [29]. These models provide the surrogate ground truth per patient, contrast, subcortical region, algorithm and sex (*s*) *Yicras*. In parallel, the *CentileBrain* model [18] estimates the subcortical volume for a healthy subject depending on age and sex, *Ŷirs*, using per-patient age and sex covariates.

**Fig 3.**
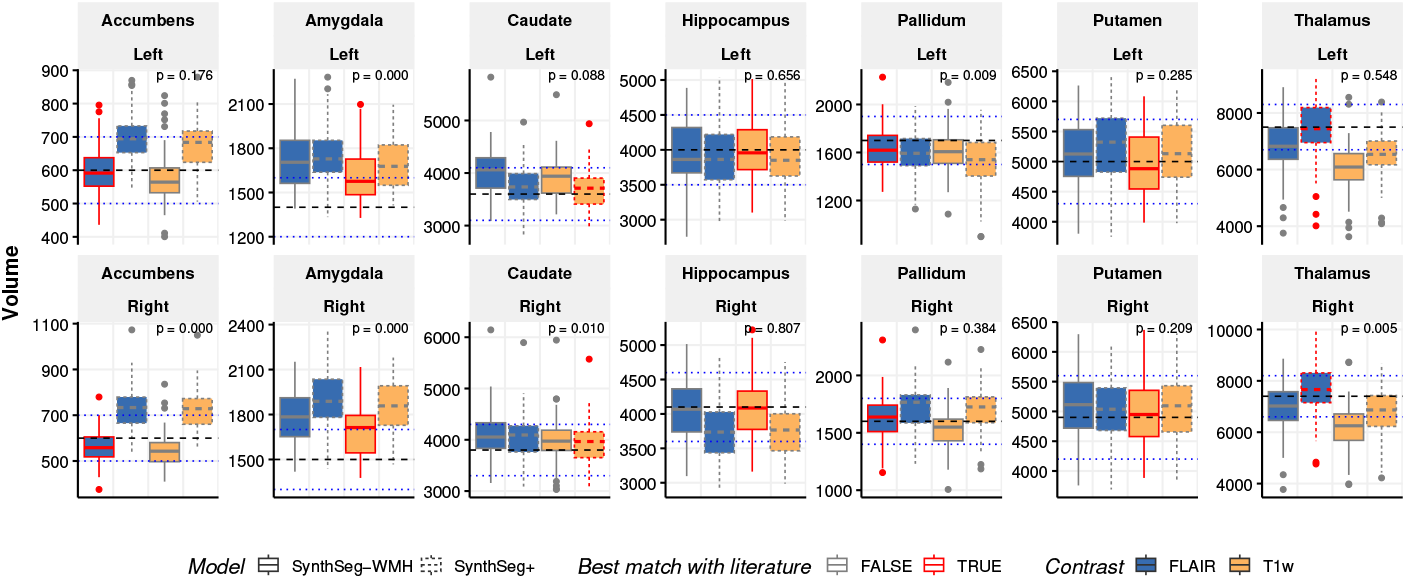
Average Estimation and Comparison with Literature-Reported Subcortical Volumes: Boxplots show the distribution of the subcortical volumes given by each model and contrast. Horizontal lines represent the literature average values for each subcortical region along with their corresponding confidence intervals [37]. Red-bordered box-plots highlight the distributions that are closest to the reference values, distinguished by the highest p-value (shown in the upper-right corners) of the four comparisons from the univariate Kolmogorov-Smirnov test.

### Subcortical Volume Longitudinal Trajectory

In Figure 4, for the sake of clarity and conciseness, we present only the results for the FLAIR contrast, as there were no significant differences for most regions except the thalamus.

**Fig 4.**
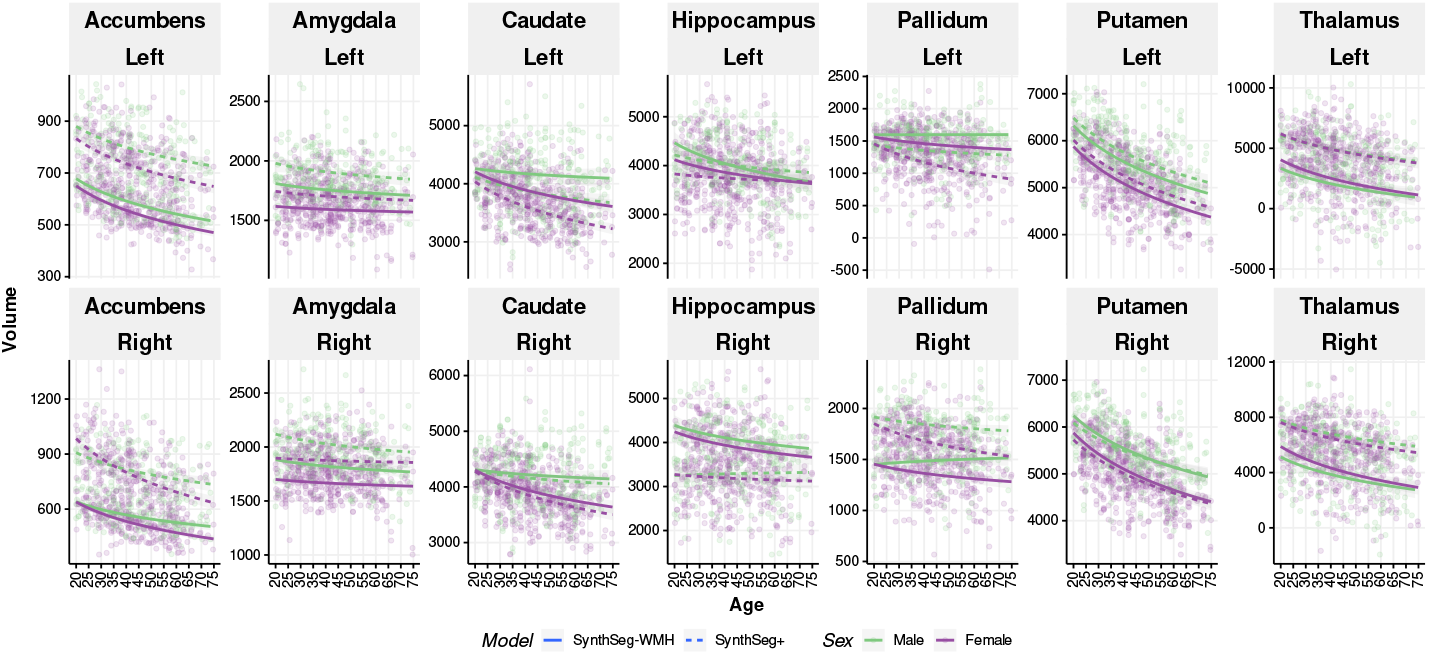
Multivariate Factorial Polynomial Regression (MFPR) [38] of each subcortical volume against the age in 326 MS patients, stratified by sex and the DL model used for segmentation employing FLAIR contrast.

All regions exhibit the expected atrophy trend, independent of sex and the segmentation model used, except for the pallidum region in males. For both hemispheres of the accumbens, amygdala, and thalamus, *SynthSeg*^+^ produced significantly larger volumes (p < 0.05, paired t-test) compared to *SynthSeg-WMH*, as well as for the right hemisphere of the pallidum, which contrasts with its trend in the left hemisphere. Conversely, the RMSE with respect to the normative model estimation for these regions behaves differently: for the left thalamus, with similar tendency in the right thalamus, it reaches 2038.95*±* 112.1 for males when using *SynthSeg-WMH* with T1w, compared to just 1304 *±*109.2 for *SynthSeg*^+^. Meanwhile, the accumbens shows an RMSE of 166.96*±*10.1 for FLAIR in males when using *SynthSeg*^+^, which is almost fifty percent less than its counterpart using T1w and *SynthSeg-WMH*, similarly to the amygdala.

### Volumetric Deviations from Normative Trajectories

Figure 5 illustrates the longitudinal trajectories of subcortical volumes for MS patients compared to the expected trajectories for healthy subjects. We observed a consistent trend across sexes, showing an increasing deviation with age in the thalamus of both hemispheres independently of the segmentation algorithm used. Although not significant, a similar trend was found for the putamen. Overall, *SynthSeg*^+^ and *SynthSeg-WMH* show significant differences in deviation when stratified by sex. In general, and consistent with the trends for subcortical volume segmentation, *SynthSeg*^+^ shows more extreme values.

**Fig 5.**
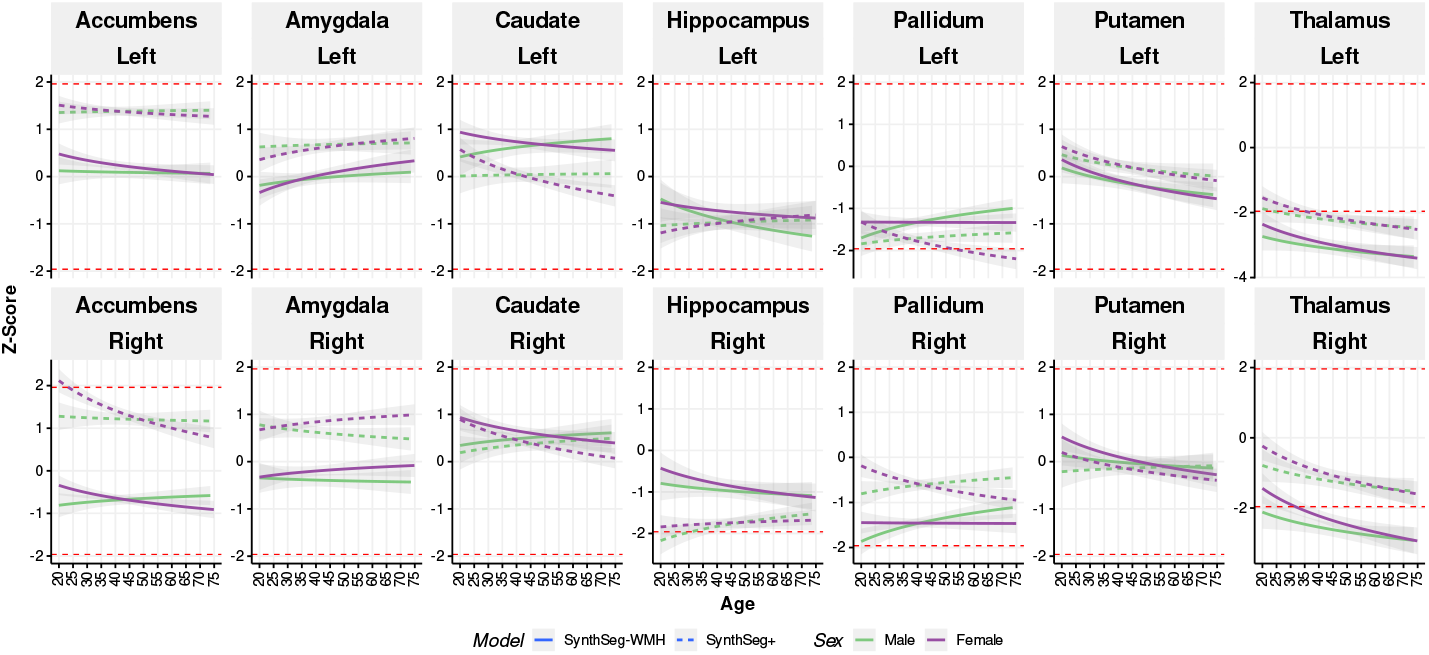
Deviation, *Z*-score, of each MS patient’s subcortical volume from the normative value for their age and sex employing the FLAIR contrast. The *Z*-score trajectory is shown as its regression against age. Red dashed lines at *±*1.96 identify the 95% confidence interval. |*Z*-score| *>* 1.96 are considered extreme deviants.

## 4 Conclusions

This pilot study explores the feasibility of normative models in identifying and quantifying brain atrophy in MS patients using state-of-the-art DL algorithms for subcortical structure segmentation, offering initial insights into disease progression. Preliminary results demonstrate that DL models can estimate subcortical volumes for MS patients. While these results are not equally relevant for all structures, it’s important to note that reference values are based on corrected automatic segmentations, often derived from T1w sequences. This could potentially bias results in favor of T1w-based methods [37], although our findings suggest this bias varies by region. The DL algorithms, in conjunction with harmonization techniques, provide results within credible ranges for all regions, as well as expectable rates of change [5, 10] as shown in Figures 4 and 5. Although direct comparisons are challenging, our findings align with literature on comparative studies for hippocampus [23], putamen [27], pallidum, and thalamus [2]. Limitations were observed in regions like the amygdala, known for its segmentation difficulties [40]. Notably, the amygdala is the only region not presenting a typical atrophy pattern (see Fig.4). Future work will incorporate lesion load analysis at specific localizations to assess its impact on volumetric measures. This will involve comparing manual lesion segmentations (not yet available for all datasets) with automatic segmentations to explore full process automation. This study confirms that FLAIR sequences can be highly relevant for adequate characterization of certain subcortical structures, such as caudate, hippocampus, and accumbens [1], paving the way for future inclusion of additional subcortical regions (currently only available through *SynthSeg*^+^). Automatic segmentation, even in the presence of WML, proves reliable, especially when using *SynthSeg-WMH*. This is evidenced by the RMSE with respect to the normative model, where *SynthSeg-WMH* provides estimates of expected deviations in MS patients with age, as well as their deviation from normativity.

In conclusion, while substantial work remains to confirm these initial findings and establish their clinical relevance, our preliminary study suggests potential subcortical volume deviations in MS patients. It primarily serves to highlight the need for further research and methodological validation. By addressing the limitations of current segmentation techniques and expanding upon this normative modeling approach, future studies can work towards more accurate DPM, improved patient stratification, and ultimately, more personalized and effective treatments for individuals with MS.

## Acknowledgments

The authors acknowledge access to facilities and expertise of the CIBM, a Swiss research center of excellence founded and supported by CHUV, UNIL, EPFL, UNIGE, HUG.

